# FOCUS2: agile and sensitive classification of metagenomics data using a reduced database

**DOI:** 10.1101/046425

**Authors:** Genivaldo Gueiros Z. Silva, Bas E. Dutilh, Robert A. Edwards

## Abstract

**Summary:** Metagenomics approaches rely on identifying the presence of organisms in the microbial community from a set of unknown DNA sequences. Sequence classification has valuable applications in multiple important areas of medical and environmental research. Here we introduce FOCUS2, an update of the previously published computational method FOCUS. FOCUS2 was tested with 10 simulated and 543 real metagenomes demonstrating that the program is more sensitive, faster, and more computationally efficient than existing methods.

**Availability:** The Python implementation is freely available at https://edwards.sdsu.edu/FOCUS2.

**Supplementary information:** available at Bioinformatics online.

## 1 INTRODUCTION

Prokaryotes are more abundant than any other group of organisms (Whitman et al., 1998), and characterizing their community composition and function is fundamental to medical, microbiological, and ecological investigations. In many environments a majority of the microbial community members cannot be cultured due to lack of information on how to grow the organisms or because some organisms only grow in groups or communities. This bias can be circumventing with metagenomics, making it a powerful tool for studying these communities. To understand the real diversity present in microbial communities in the wild, researchers use next-generation sequencing (NGS) and metagenomic profiling. These high-throughput technologies are promising for biological research avenues in which rapid turnaround time and large-scale characterization is advantageous. For example, metagenomic profiling can discriminate taxonomic profiles of microbes associated with human (Consortium, 2012a) and global ocean microbiomes (Sunagawa *et al*., 2015).

Metagenomics uses high throughput sequencing, a fast and cheap sequencing method provided by recent NGS technologies.

An alternative to metagenomes is metabarcoding, where DNA from a specific marker gene, such as 16s rRNA, is sequenced in order to understand the whole community. This approach only characterizes the taxonomic profile of community and it ignores the functional profile. The lack of functional characterization by metabarcoding makes metagenomics a more complete approach (see Figure 1 for a representation of the whole pipeline), with complex bioinformatic attempts to bridge the gap between barcoding and functional information [e.g., Tax4fun (Aßhauer *et al*., 2015) and PICRUSt (Langille *et al*., 2013)].

**Fig. 1.**
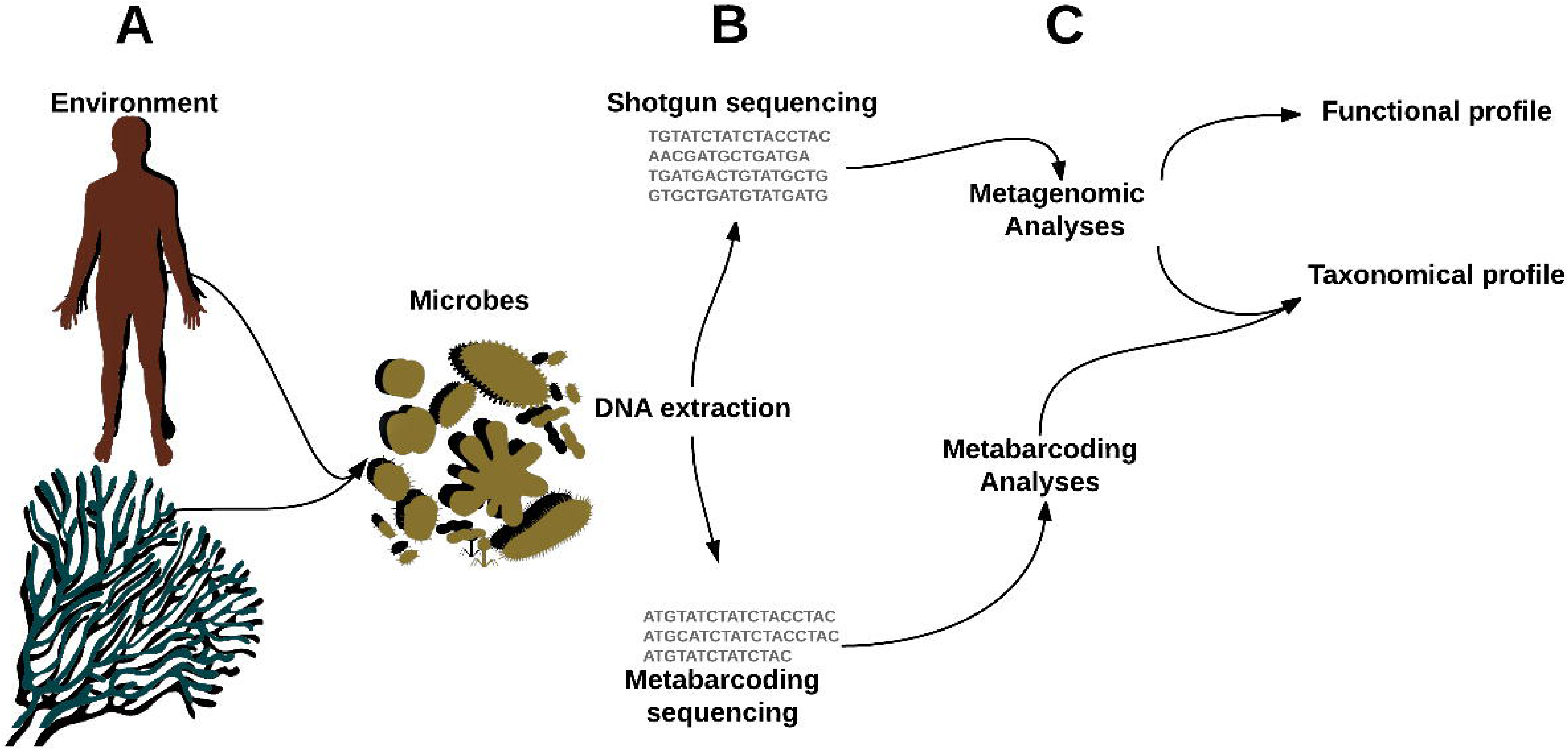
Environmental analysis: A) A sample is collected from the environment; B) DNA is extracted and sequenced which produces sequences from many species; and C) Depending on the study,only some sequences from the samples are targeted such as 16S rRNA for taxonomy; metagenomics is done when a functional understanding of the community is needed.

Large metagenomic datasets are increasingly being generated due to declining sequencing cost and increasing access to sequencing platforms. However, many of the available tools do not scale well with increasing data volumes, precluding timely analysis or capitalizing on the depth of these datasets.

We previously published FOCUS (Silva *et al*., 2014), an accurate computational method using non-negative least squares and 7-mers to profile large metagenomic datasets in seconds. Here we propose an update to FOCUS, named FOCUS2, which guarantees more accurate taxonomic assignment of metagenomic datasets via homology to a reduced database. FOCUS2 was validated using simulated data and two large datasets from the Human Microbiome Project (HMP) (Consortium, 2012a) and Tara global ocean expedition (Sunagawa *et al*., 2015). Our approach was more sensitive, agile, and computationally efficient when compared to existing tools.

## 2 METHODS

The FOCUS2 workflow is presented in Fig. 2 and described below:

1. Resample 80% (default; see Suppl. Methods) of the sequences in the input.
2. Profile the resampled sequences via FOCUS using the PATRIC database (Wattam et al., 2013). Repeat steps 1 and 2 “n” times (n = 100 by default).
3. Create a reduced database containing genomes present in at least 90% (default) of the profiles.
4. Align input sequences against the reduced database using blastn/HS-blastn (Chen *et al*., 2015a) (Aligner choice discussed in “Aligner choice” below).
5. Write the classification for each sequence identified.

**Fig. 2.**
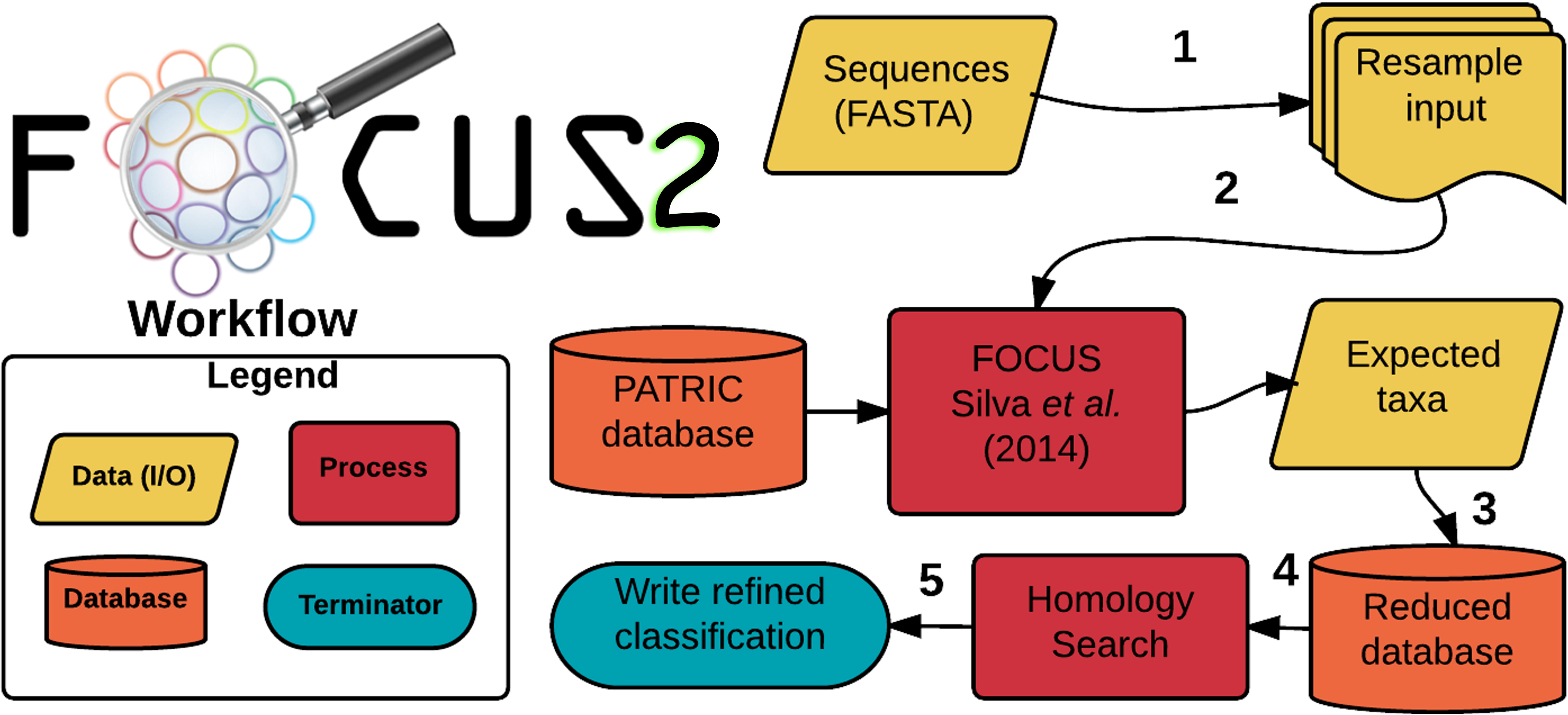
Workflow of the FOCUS2 program.

### 2.1 Aligner choice

Blastn is an available aligner choice for in FOCUS2. However, HS-blastn (Chen et al., 2015b) is the default aligner because it is up to 22x faster than blastn with the same sensitivity. Any other aligner, which generates a tabular output, can be easily integrated into the FOCUS2 pipeline.

### 2.2 Reference dataset

FOCUS2 expands the FOCUS database by using over 33,000 complete and draft genomes (~12 times more genomes than FOCUS) from the PATRIC platform. *K*-mer counting and normalization of the database were done as for FOCUS.

### 2.3 Resampling of the data via Monte Carlo simulation

We implemented an optional resampling strategy on step 1) of the FOCUS2 pipeline using Monte Carlo simulation to assess the confidence that organisms identified were present in input samples. 80% of the reads were randomly resampled 10 times, and the taxa frequencies recalculated. The species present in at least 80% of the profiles are considered robust/reliable taxa, and they are used to create the reduced database for step 4) of the tool pipeline.

### 2.4 Simulated and real testing set

FOCUS2 was evaluated with ten simulated big datasets (total of 100 million reads) composed of the same taxa in the “HiSeq” and “MiSeq” datasets used by (Ounit *et al*., 2015). However, we recreated them using BEAR (Johnson et al., 2014) with the same number of sequences per sample, same taxa abundance, but with different sequences lengths (100, 250, 500, 750, and 1,000 bp). In addition, 300 real dataset from the HMP from 15 sites and 243 samples from the Tara project (Suppl. Table 1) were selected as test sets.

### 2.5 Sensitivity and precision FOCUS2 analysis

Analyses of simulated data were evaluated by sensitivity, the ratio between the number of correct assignments by FOCUS2 and the total number of sequences in the sample, and precision, the ratio between the number of correct assignments by FOCUS2 and the total number of classified sequences by FOCUS2.

### 2.6 Memory usage and speed: FOCUS2 vs CLARK

In order to compare speeds between FOCUS2 and CLARK version 1.2.2-b (Ounit et al., 2015), we analyzed 100 metagenomes from the HMP (Consortium, 2012b) using one thread: FOCUS2 was set to use the HS-blastn aligner with no resampling, and CLARK was set to run in the default and full mode [high confidence assignments (i.e, confidence score >= 0.75, gamma score >= 0.03)] with k-mers of 21. FOCUS2 was able to classify 841,742 reads per minute, while CLARK classified 1,113,658 reads per minute in full mode and 3,492,330 in default mode. However, FOCUS2 used ~2GB of RAM while CLARK on the full mode required ~70.7 GB, and ~47 GB on its default mode. CLARK has a light version (CLARK-l), which requires ~4GB of RAM; however, it has sensitivity of only ~60 % (http://clark.cs.ucr.edu). FOCUS is ~10x faster than CLARK for databases of comparable size if we normalize the runtime by the database size.

## 3 RESULTS AND DISCUSSION

All the tools were run using the same database on a server with 24 processors x 6 cores Intel(R) Xeon (R) CPU @2.67 GHz and 189 GB RAM.

### 3.1 Comparision of FOCUS2 and other tools

Ten simulated metagenomes were analyzed using FOCUS2 and the results were compared to blastn best hit assignments (E-value 10^-5^, min. 60% identity, and min. alignment length 15 amino acids). Fig. 3a shows that FOCUS2 is more sensitive and precise when compared to blastn for genus-level binning, and much more sensitive and precise than blastn in the species level (Fig. 3b).

**Fig. 3.**
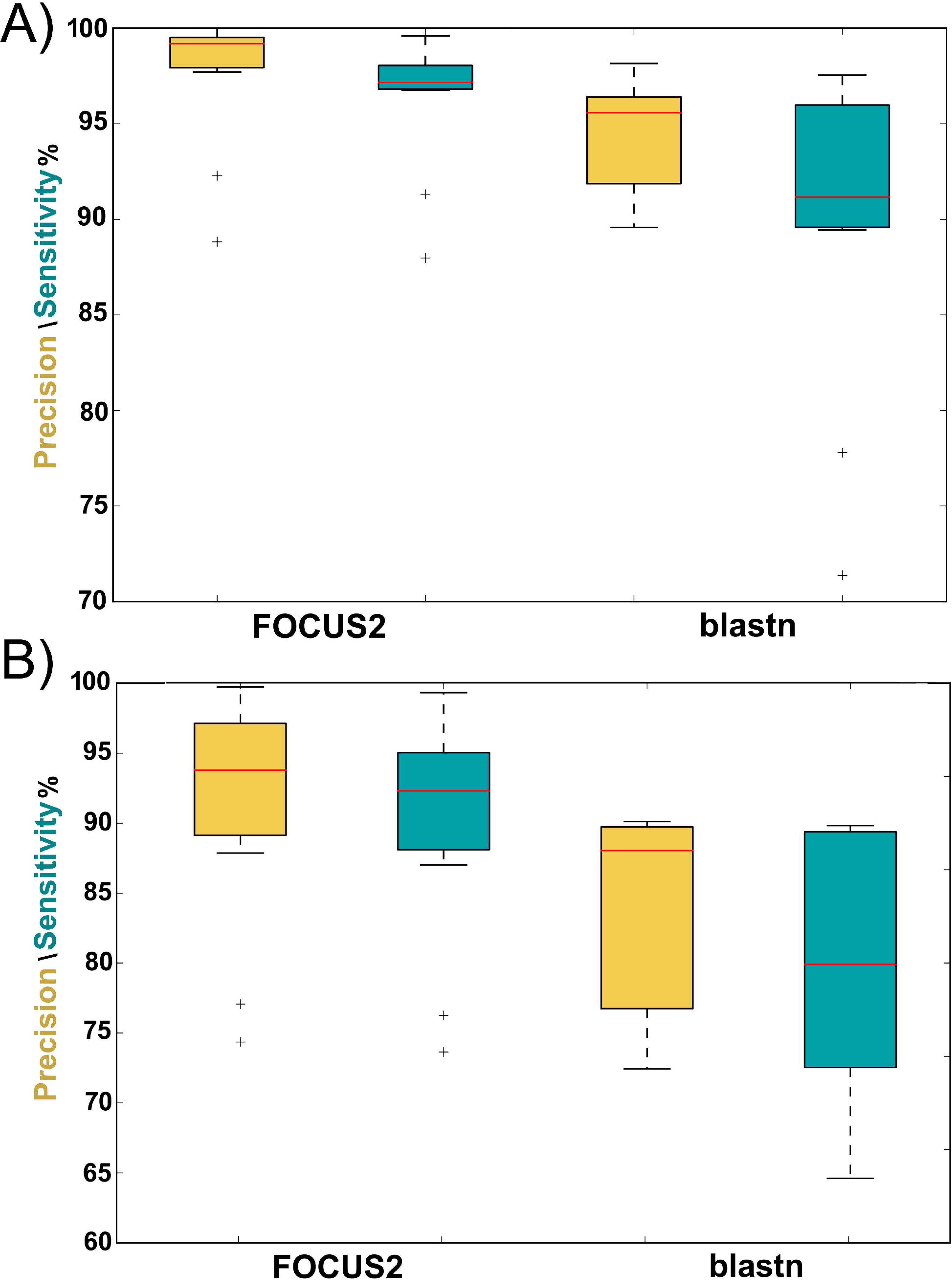
Box plots displaying the percent precision (yellow) and sensitivity (turquoise) of FOCUS2 and blastn binning assignments in the genus (A) and species (B) level for 10 simulated metagenomes.

For the 543 real metagenomes from the HMP and the Tara expedition, we analyzed the data using FOCUS2, FOCUS, CLARK (Ounit *et al*., 2015) in default [CLARK (D)] and full mode [CLARK (F)] and displayed the results a normalized heat-map of distance matrices Fig. 4.

**Fig. 4.**
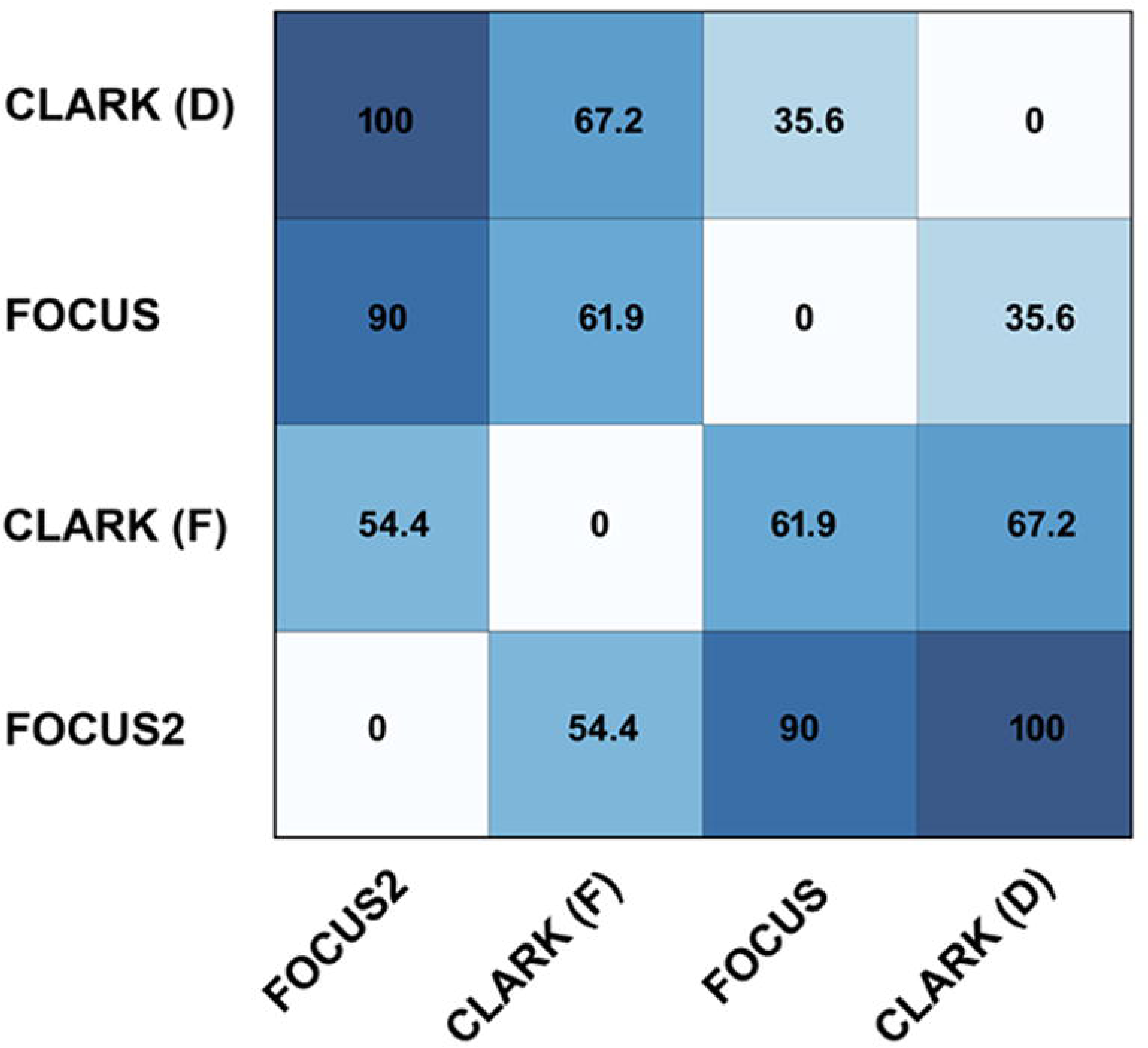
Normalized heat-map generated from distance matrices of FOCUS(2) and CLARK (D/F).

To compare the HMP and TARA datasets against the full PATRIC dataset, FOCUS2 required ~6GB of RAM. No other metagenomic profiling tool could be used to analyze these metagenomes with the full PATRIC database. In order to compare FOCUS2 to any other tool, we use the state-of-the-art binning tool, CLARK, but had to used a reduced dataset of ~2,800 genomes (the dataset we used in the original FOCUS paper). Even with this reduced dataset, CLARK required ~70.7 GB of RAM. FOCUS2 is ~10x faster than CLARK for databases of comparable size (see Methods). A comparison of FOCUS(2) to CLARK (D/F; heat-map on Fig. 4) shows that the FOCUS2 profile is closer to CLARK (F) (and vice-versa) and that FOCUS is closer to CLARK (D), which suggests that FOCUS2 is as highly sensitive as CLARK (F). The same is suggested by the hierarchical clusters on Fig. 5 and 6.

**Fig. 5.**
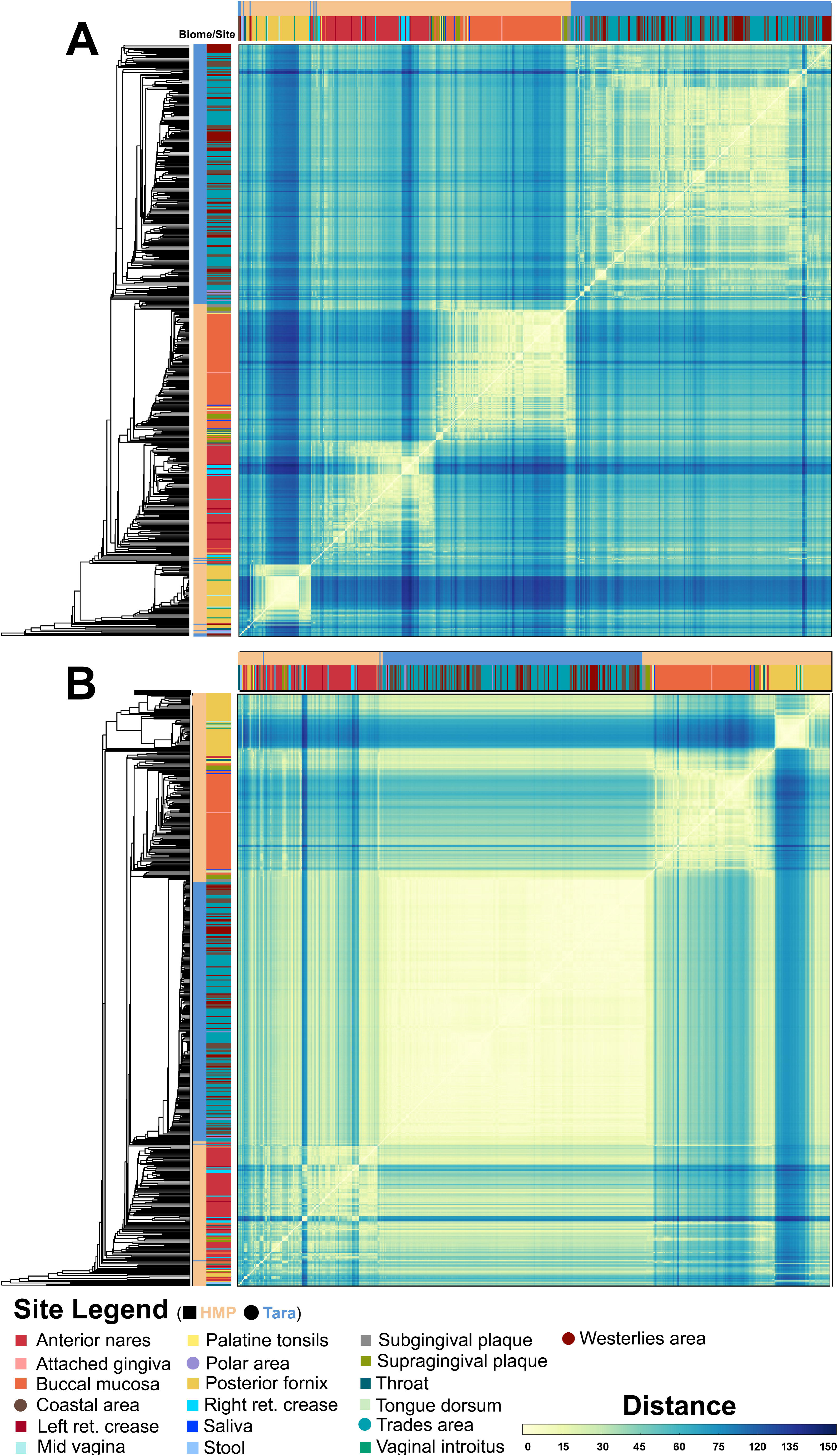
Hierarchical clustering of genus level taxonomic annotation performed using FOCUS2 (A) and FOCUS (B) on 543 metagenomes from the HMP (squares) and Tara ocean expedition (circles) The color bars fringing the similarity plot represent the human (HMP; tan) and ocean (Tara expedition; blue) biomes sampled and the 19 sites on those biomes.

**Fig. 6.**
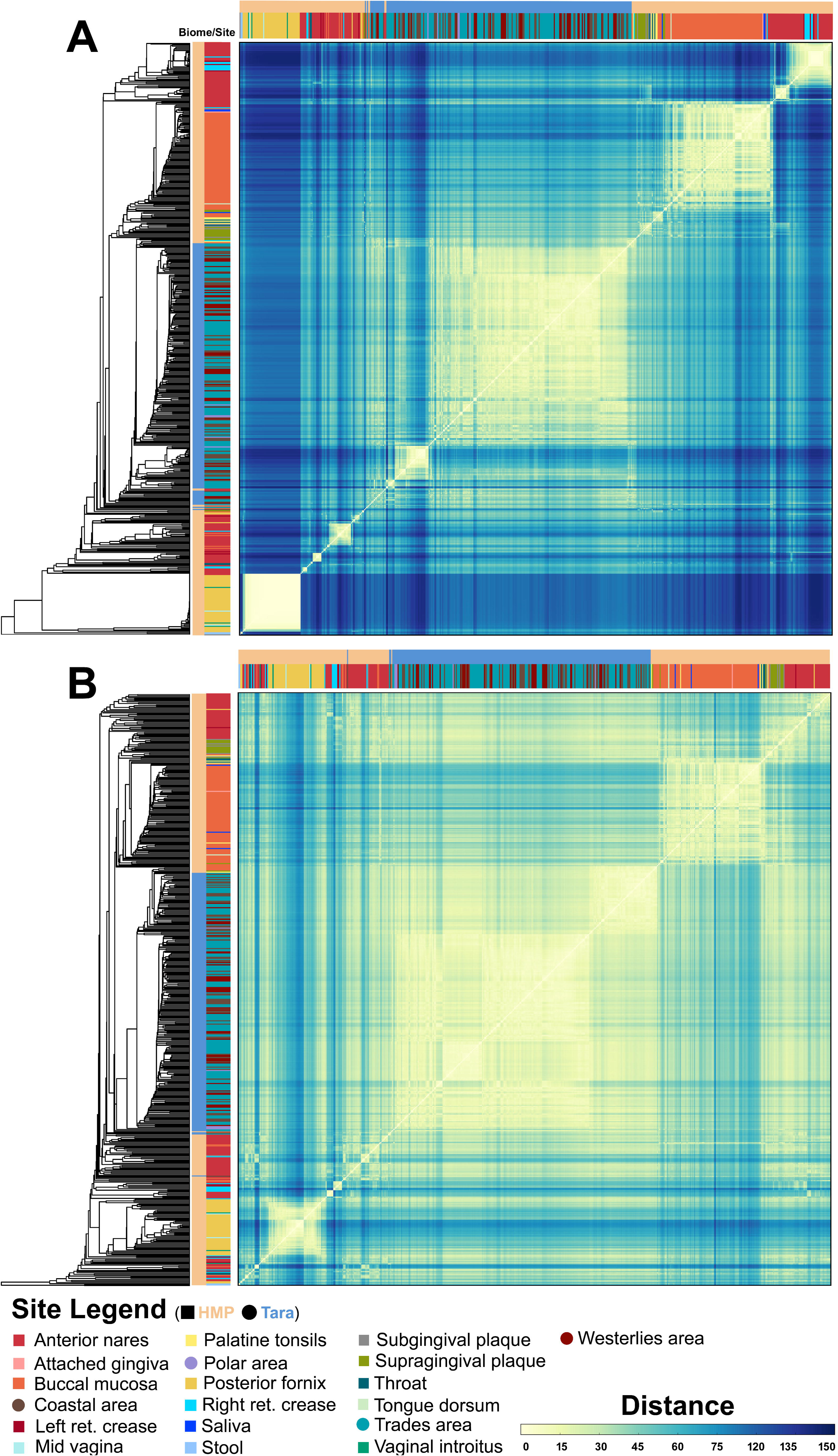
Hierarchical clustering of genus level taxonomic annotation performed using CLARK (A) in full mode considering only high confidence assignments; and (B), in default mode on 543 metagenomes from the HMP and Tara ocean expedition. The color bars fringing the similarity plot represent the human (HMP; tan, squares) and ocean (Tara expedition; blue, circles) biomes sampled and the 19 sites on those biomes.

### 3.2 Final considerations

FOCUS2 is a sensitive and fast solution to bin metagenomic samples. It first runs FOCUS to predict the taxa in the sample taxa and refines the profiling using a fast aligner with a reduced version of the PATRIC database created on the fly. The PATRIC database opens new horizons in the metagenomics binning world because it is over 12x bigger than previous databases and brings many new taxa into classification. The speed, sensitivity, and precision of FOCUS2 positions metagenomics to capitalize on expanding databases and ask novel interdisciplinary questions currently beyond reach.

## ACKNOWLEDGMENTS

GGZS thanks Ben Knowles and Kathryn Furby for the comments on the manuscript. Dr Elizabeth Dinsdale and Daniel Cuevas for the discussion in the manuscript.

